# Fuzzy ripple artifact in high resolution fMRI: identification, cause, and mitigation

**DOI:** 10.1101/2024.09.04.611294

**Authors:** Renzo Huber, Rüdiger Stirnberg, A Tyler Morgan, David A Feinberg, Philipp Ehses, Lasse Knudsen, Omer Faruk Gulban, Kenshu Koiso, Stephanie Swegle, Isabel Gephart, Susan G Wardle, Andrew Persichetti, Alexander JS Beckett, Tony Stöcker, Nicolas Boulant, Benedikt A Poser, Peter Bandettini

## Abstract

**Purpose:** High resolution fMRI is a rapidly growing research field focused on capturing functional signal changes across cortical layers. However, the data acquisition is limited by low spatial frequency EPI artifacts; termed here as Fuzzy Ripples. These artifacts limit the practical applicability of acquisition protocols with higher spatial resolution, faster acquisition speed, and they challenge imaging in lower brain areas.

**Methods:** We characterize Fuzzy Ripple artifacts across commonly used sequences and distinguish them from conventional EPI Nyquist ghosts, off-resonance effects, and GRAPPA artifacts. To investigate their origin, we employ dual polarity readouts.

**Results:** Our findings indicate that Fuzzy Ripples are primarily caused by readout-specific imperfections in k-space trajectories, which can be exacerbated by inductive coupling between third-order shims and readout gradients. We also find that these artifacts can be mitigated through complex-valued averaging of dual polarity EPI or by disconnecting the third-order shim coils.

**Conclusion:** The proposed mitigation strategies allow overcoming current limitations in layer-fMRI protocols:

1. Achieving resolutions beyond 0.8mm is feasible, and even at 3T, we achieved 0.53mm voxel functional connectivity mapping.
2. Sub-millimeter sampling acceleration can be increased to allow sub-second TRs and laminar whole brain protocols with up to GRAPPA 8.
3. Sub-millimeter fMRI is achievable in lower brain areas, including the cerebellum.

## 1.) Introduction

### 1.1) Layer-fMRI and its acquisition challenges

Layer-fMRI has significant potential for investigating neural information flow within and across brain systems. Knowing at which cortical layer neural activity occurs allows neuroscientists to determine whether activation modulations are driven by feed-forward or feedback input, and whether neural circuits are involved in output vs. input processes.

However, traditional layer-fMRI data collection at submillimeter resolutions faces limitations due to EPI artifacts. Specifically, in high-resolution protocols with low bandwidths, EPI ghosts arise from inconsistencies between odd and even echoes. While conventional Nyquist ghosting is addressed through common two-parameter phase correction method^1–3^, layer-fMRI is also affected by additional higher-order effects:

- Imperfections in gradient waveforms increase with higher gradient amplitude and slew rates. This is particularly challenging for EPI image quality when large ramp sampling factors and low bandwidths are used in layer-fMRI protocols.
- The long echo train length in high-resolution protocols allows phase inconsistencies to accumulate during the acquisition of large imaging matrices, especially in brain areas with stronger B0 inhomogeneities.
- Parallel imaging, which is crucial for efficient layer-fMRI, relies on “known” aliasing patterns, meaning even small residual EPI ghosting and phase errors are amplified with GRAPPA/SENSE.

Due to these challenges, conventional layer-fMRI acquisition protocols are often conservatively designed, with limited resolutions of approximately 0.8mm and TR values around 3 seconds. This resolution is barely sufficient to reveal differences in neuronal laminar activation. Additionally, field of view prescriptions are typically restricted to cover only the outer cortical areas of the upper cortex. This is because:

- Many lower brain areas are located far from the receive RF-coil elements, leading to high g-factors and low SNR.
- Lower brain areas experience stronger B0 inhomogeneities, resulting in more severe EPI artifacts.
- Lower brain areas require large matrix sizes for acquisition, but shorter T2* values lead to increased signal decay and reduced SNR.

Among the 265 human layer-fMRI papers published between 1997 and 2024, fewer than five focus on low ventral brain structures (source https://layerfmri.com/papers).

The goal of this study is to:

1. Characterize a significant limitation of high resolution fMRI, referred to as “*Fuzzy Ripple* artifacts”, that prevent researchers from advancing to Cartesian EPI protocols with smaller voxels, shorter TRs, and coverage of lower brain areas.
2. Identify the primary cause of this artifact: EPI trajectory imperfections in ramp-sampling readouts due. Characterizing the exacerbation of Fuzzy Ripples with a number of system imperfections: Including (i) gradient imperfections caused by inductive coupling between third-order shim coils and readout gradients, (ii) interactions of imperfect B0 shim (typical in lower brain areas) with trajectory imperfections, (iii) interactions of aggressive GRAPAP accelerations with trajectory imperfections.
3. Implement and test a potential mitigation strategy: complex-valued averaging of dual-polarity EPI readouts, and unplugging the third order shims.
4. Empirically evaluate whether this mitigation strategy enables neuroimagers to surpass the current resolution limits of conventional layer-fMRI protocols, with respect to TR, voxel size, and imaging of lower brain areas. Our focus will be on practical aspects in the most commonly used sub-millimeter fMRI sequences, specifically the 3D-EPI from DZNE^4^ with and without VASO^5^ and, to some extent, the Multi-Band C^2^P from CMRR^6^ on SIEMENS (Siemens Healthcare, Erlangen, Germany) 7T scanners equipped with the SC72 gradient coil which incorporates third order shimming capabilities.

### 1.2) Fuzzy Ripple artifacts are relevant for the field of layer-fMRI

Previously acquired EPI brain images, shown in Figs. 1–2, highlight the types of imaging artifacts that pose limitations for high-resolution fMRI. We refer to these artifacts as “Fuzzy Ripples.”

**Fig 1:**
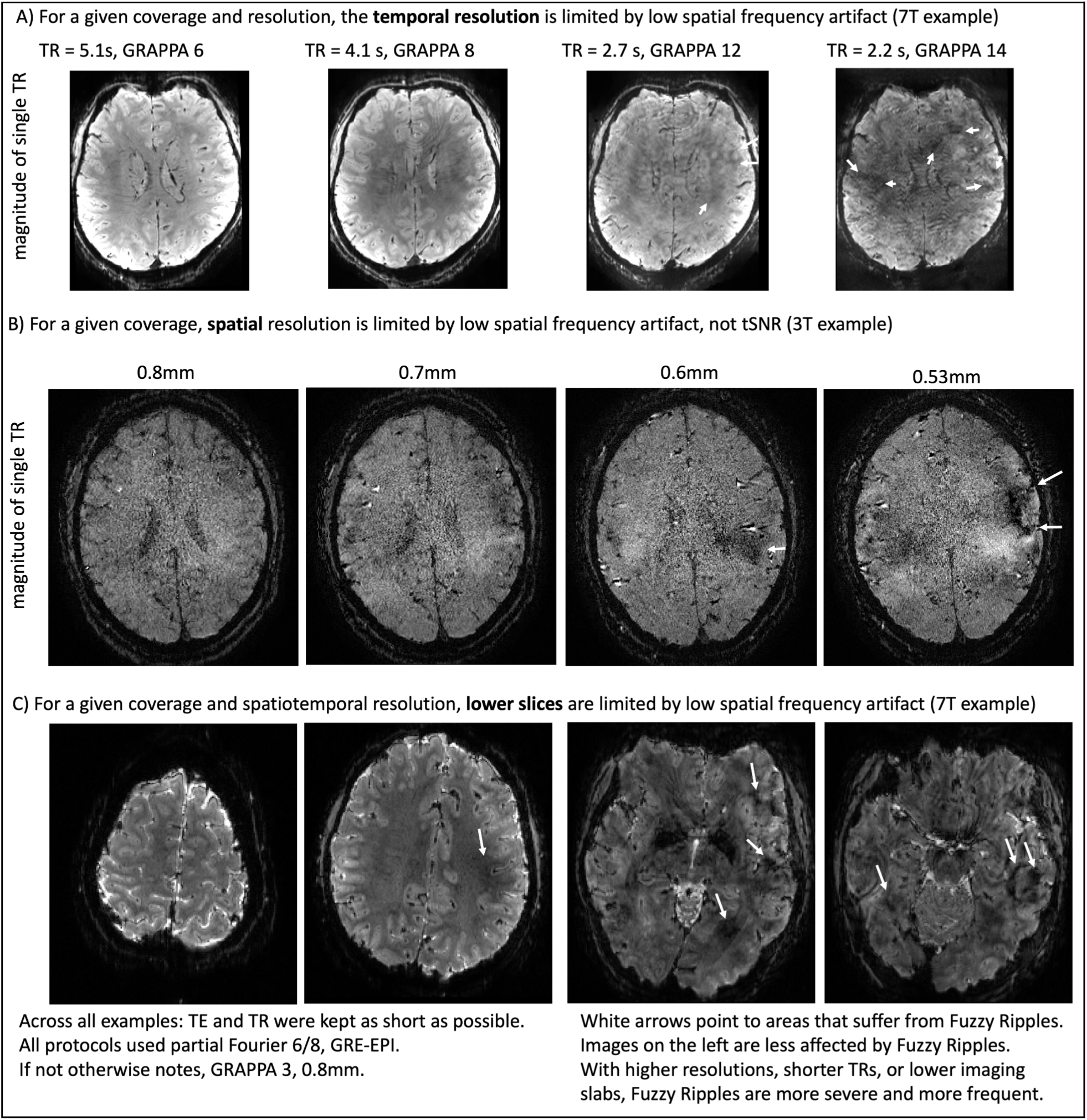
Fuzzy Ripples are the reason why layer-fMRI is confined to conventional protocols. Fuzzy Ripples are the primary reason why layer-fMRI is restricted to conventional protocols. Standard layer-fMRI protocols are generally limited to 0.8mm resolution with TRs of several seconds, focusing on upper cortical brain areas. These limitations cannot be surpassed because, with more ambitious acquisition protocols. Fuzzy Ripple artifacts become too strong and too frequent. Panel **A)** exemplifies issues of pushing TR. Panel **B)** exemplifies issues of pushing resolutions beyond 0.8mm. Panel **C)** exemplifies issues of pushing protocols towards lower brain areas. Images from Panels A and B are taken from Koiso 2023 and Huber 2023, respectively. They refer to 3D-EPI readouts with planar EPI trajectories.

**Fig 2:**
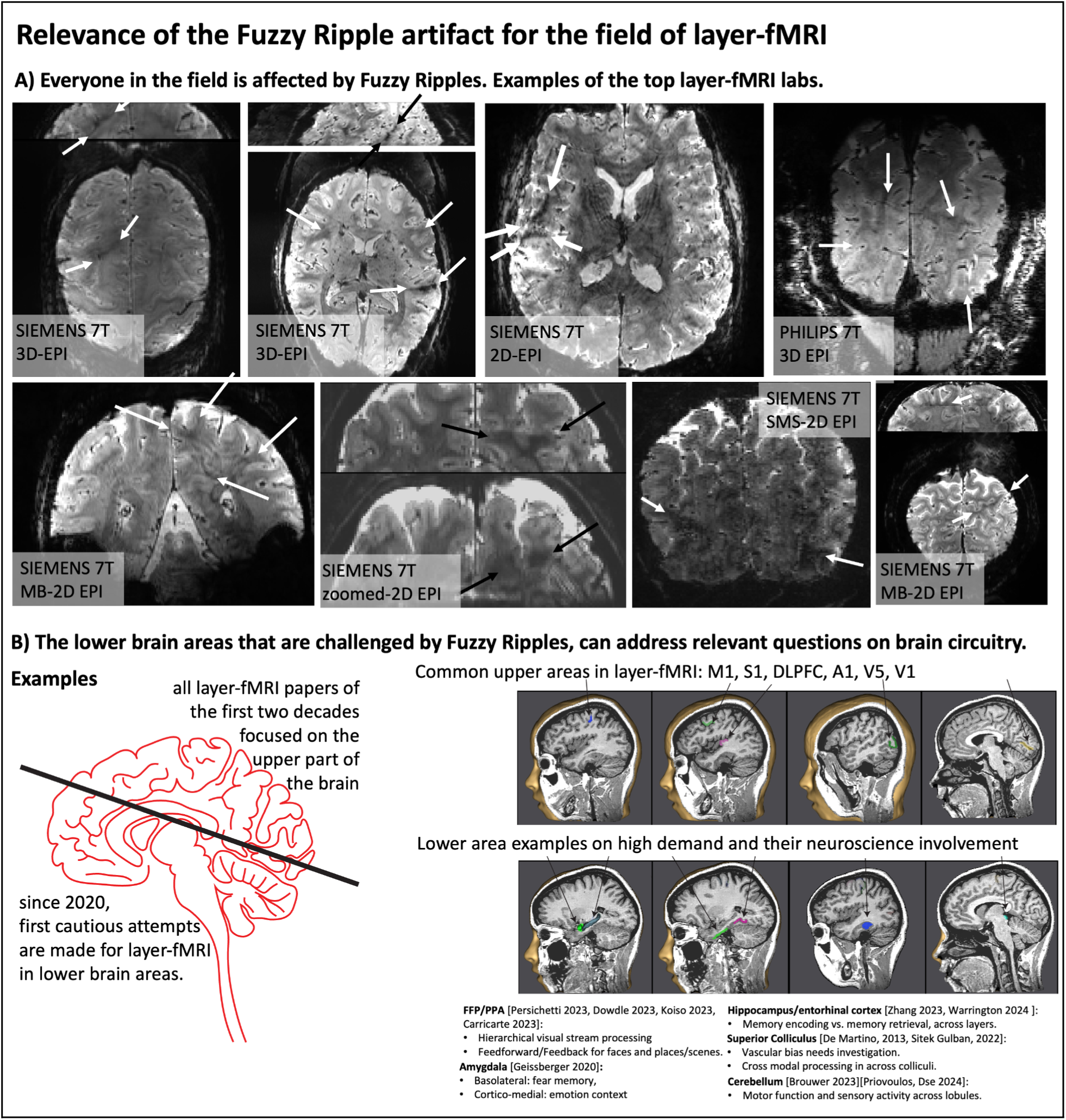
Fuzzy Ripples and their impact on layer-fMRI research. **Panel A). The widespread effect of fuzzy ripples:** Representative EPI images from among top 10 layer-fMRI labs (based on number of publications on www.layerfmri.com/papers): Maastricht, Nijmegen/Essen, CMRR, NIH, MGH, Amsterdam, Leipzig, Cambridge. Note Utrecht/Tübingen are excluded, as despite being among the top 10 layer-fMRI labs, none of their papers include publicly available layer-fMRI EPI data. **Panel B). The Necessity of High-Resolution fMRI in Lower Brain Areas**. High-resolution fMRI in lower brain areas is crucial for addressing key open questions about human brain circuitry. While most layer-fMRI studies have focused on upper brain regions, lower brain areas are equally important for fundamental neuroscience research. Examples of relevant research topics include: feedforward vs. feedback processing in layers of ventral cortical areas like the FFA/PPA, differential processing in mesoscale subnuclei of the amygdala, laminar differentiation of memory encoding and retrieval in the hippocampus and entorhinal cortex, multi-modal sensory integration across the colliculi, and mesoscale representations of body parts in the fine-scale lobules of the cerebellum. The underlay images in Fig. 2 were generated using the Brain Tutor app for Android by Brain Innovations (Rainer Goebel).

Fig. 1 illustrates how the severity of these artifacts increases when conventional layer-fMRI protocols—featuring 0.8mm resolution, TRs of several seconds, and imaging of upper brain areas—are pushed to shorter TRs, smaller voxels, and lower brain regions. While the conventional protocols (left column) may already show faint artifacts, the right columns demonstrate how these artifacts can become so severe that the data is rendered unusable. In the right column, it can be seen that these artifacts are the primary source of noise, more so than the thermal noise, which appears as a salt-and-pepper graininess in the images.

Although these artifacts are most pronounced in more aggressive protocols, they are also present in conventional protocols (0.8mm, upper brain, few sec. TR), although usually at a lower magnitude that doesn’t necessarily render the datasets unusable. Instead, they limit scanner operators’ ability to achieve higher spatiotemporal resolutions across different brain areas in the first place. Figure 2A shows representative high-resolution EPI data from the top ten labs with the most published human layer-fMRI papers. Without exception, Fuzzy Ripples are visible in all of them, underscoring the significance of this artifact for the entire research field.

Across all the EPI brain images shown in panels of Figs. 1–2, artifacts share common characteristics. Specifically, they exhibit local signal intensity deviations at low spatial frequencies, which can be described as fuzzy clouds of brighter or darker signals. The darker Fuzzy Ripples are generally more noticeable to the naked eye. While these artifacts may contain wave-like warble patterns at higher spatial frequencies, their overall appearance is usually blurry. The size of the artifacts typically ranges from 5% to 15% of the field of view (FOV). In this study, our goal is to identify the origins of Fuzzy Ripples and explore strategies for mitigating them.

## 2.) Theory

### 2.1) readout direction specific eddy currents in ramp-sampled EPI can result in odd-even ghosts at low spatial frequencies

In conventional high resolution Cartesian EPI, as used in common 2D/Multiband-EPI and 3D-EPI, readout gradients are driven at their maximum allowed amplitude and slew rate. This operation pushes the gradients beyond the regime of vendor-provided optimal eddy current compensation calibration^7^.

Furthermore, the design of conventional body gradients is not optimized for high-resolution, encoding-limited head EPI. As a result, the trapezoidal gradient pulse shapes heavily depend on data sampling along the slopes, with ramp sampling fractions reaching up to 78% of the entire duration of the read pulses. This is outside the range of conventional vendor-provided EPI ghost correction methods, which typically assume that gradient delays can be corrected with line-wise two-parameter phase correction^1–3^. With such high ramp sampling ratios, gradient delays introduce readout direction-dependent phase offsets that vary significantly throughout the gradient pulse evolution.

Figure 3A illustrates the gradient shape and trajectory imperfections for the protocol shown in Fig. 1A^8^. It can be seen that the strongest gradient imperfections occur at the corners of the trapezoidal pulse shapes, which cannot be adequately corrected by conventional vendor-provided Nyquist ghost correction strategies. Instead, a global phase correction scheme may lead to residual phase errors that are distributed differently across representations of low and high spatial frequencies (Fig. 3B). In this study, we hypothesize that these mechanisms may partially contribute to the Fuzzy Ripple artifacts.

**Fig 3:**
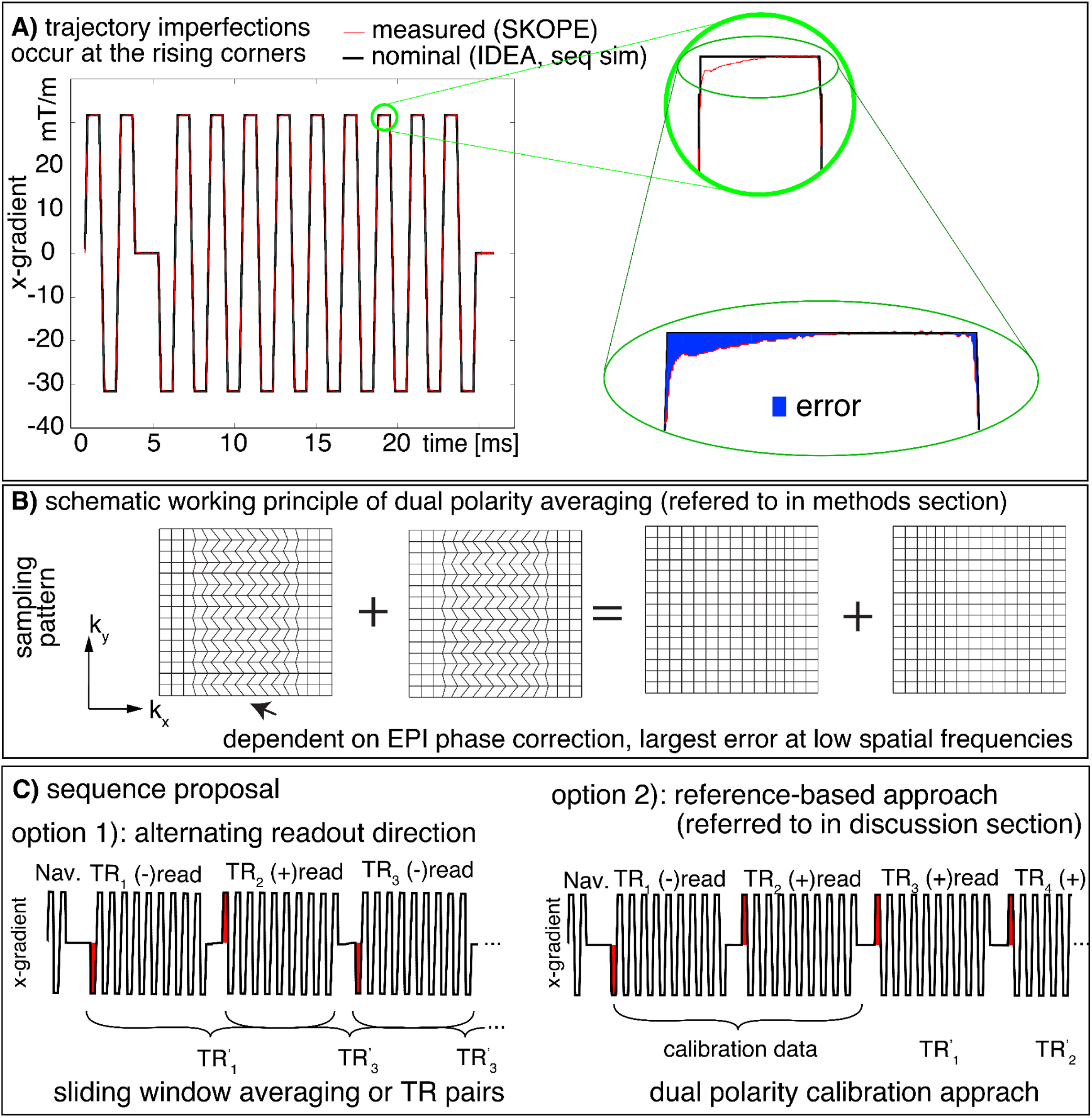
Concept of Fuzzy ripples as EPI odd-even delays with ramp sampling with readout (here k_x_)-specific phase errors. **A)** Gradient imperfections in high resolution EPI are most prominent at the corners of trapezoids,including both the rising and falling edges. The nominal EPI trajectory is derived from SIEMENS IDEA simulations, while the measured trajectory is obtained using SKOPE on a standard SIEMENS 7T MAGNETOM with third-order shimming. **B)** Despite phase correction in the image reconstruction process, residual imperfections persist in the part of k-space that encodes lower spatial frequencies. These residual errors, shown as deviations in k_x_, manifest as irregular k-space grids for odd and even lines. Such odd-even errors are expected to produce EPI ghosting artifacts in low spatial frequencies. **B-C)** This study introduces a strategy to address Fuzzy Ripple artifacts by employing a dual-polarity EPI approach that alternates the read direction every other TR (see methods section). EPI images with opposite read directions are anticipated to produce ghosts with opposite phases. Option 2 refers to an alternative future implementation that is referred to in the discussion section.

### 2.1) Potential mitigation strategies of Fuzzy Ripples

#### 2.1.1) Dual-polarity averaging

Combining dual-polarity readouts to separate echoes of opposite polarities and performing complex-valued averaging on their respective images has been proposed as a method to reduce off-resonance-induced ‘edge-ghosts’ (high spatial-frequency ghosts) in EPI-based fMRI acquisitions^9–13^ and others. The principle of this approach is illustrated in Fig. 3B-C. Reversed readout polarities are expected to completely invert the k-space shift pattern, meaning that the resulting EPI ghosts should have opposite phases for each readout polarity and can be effectively canceled through complex-valued averaging. In this study, we explored whether this strategy can also address low-frequency ripple artifacts.

#### 2.1.2) Mitigation of eddy currents by minimizing inductive coupling between gradient and third-order shim coils

Recent observations by Boulant et al. (2024) indicate that third-order shim coils influence gradient-magnet interactions. Due to the shared geometric symmetry of the third-order shim coils and the gradients, inductive coupling can occur, leading to compromised gradient impulse response functions at certain frequencies. We investigated the impact of third-order shims on Fuzzy Ripples and assessed whether disconnecting the third-order shims could serve as a mitigation strategy.

## 3.) Methods

We scanned 29 participants as part of this study, all of whom provided informed consent. Twenty-four participants were scanned at NIH, while the remaining participants were scanned during the pilot phases at the University of Maastricht and the University of California, Berkeley, with each institution’s local IRB approval, in accordance with the Declaration of Helsinki. Complete scanning protocols and sequence parameters are available here: https://github.com/layerfMRI/Sequence_Github/

### 3.1) Experiments to exemplify limits of sampling speed at conventional resolutions

We estimated the limitations of TR in three sessions using a SIEMENS classic Magnetom 7T with SC72 gradient sets and 0.8 mm isotropic resolution 3D-EPI. This protocol followed previously established guidelines^8^. The full protocol parameters can be found here: https://github.com/layerfMRI/Sequence_Github/blob/master/Whole_brain_layers/20211012_KEN.pdf. We also assessed EPI trajectory imperfections using a Skope (Skope MRT, Zürich, Switzerland) clip-on field camera from a separate scan. Results from these three sessions are shown in Fig. 1A and Fig. 2.

### 3.2) Experiments to exemplify resolution limits at 3T

Most relevant scan parameters include: SIEMENS Prisma 3T (with XR-gradient set), 3D-EPI with isotropic resolutions between 0.8 and 0.53 mm. The full list of protocol parameters is available here: https://github.com/layerfMRI/Sequence_Github/blob/master/dual-polarity/3T_inverted_res.pdf Functional activation was induced using a free movie-watching paradigm, utilizing the same 15-minute movie clips from the 7T HCP study (MOVIE1).

### 3.3) Large FOV data to compare Fuzzy Ripples with edge EPI ghosts

Two participants were scanned to compare Fuzzy Ripples with conventional edge ghosts. A large FOV of 400 mm was used to capture Fuzzy Ripple ghosts independently of the main signal. The main imaging parameters were consistent across experiments: 3D-EPI, resolution 1 mm isotropic, TE 26 ms, echo spacing 1.02 ms, 7T with SC72 gradient sets. Comparisons were made with and without: GRAPPA, ramp sampling, a good shim, and dual-polarity averaging. Full protocol details are available here: https://github.com/layerfMRI/Sequence_Github/blob/master/Terra_protocolls/Fuzzy_ripples/20230424_largeFOV.pdf

### 3.4) Auditory activation with 3rd order shim induced Fuzzy Ripples

To illustrate the impact of Fuzzy Ripples on functional time series, task-based activation experiments were conducted using the protocols mentioned above. Functional runs followed previous experiments^15^ and consisted of 14 minutes with alternating 30-second blocks of rest and auditory stimulation. Sounds were delivered using MRI-compatible ear buds from Sensimetrics Corporation (www.sens.com), with tones described as “Chipmunks from space”. Sound sample available here: https://youtu.be/TGX_Ulbv9wA?si=VuhMcQj_vDYkibhc. Four experiments were conducted with two participants each participating twice. Protocol parameters included: 0.8 mm resolution, 2D-multiband sequence from CMRR^6^, echo spacing 1.0 ms, TE 26 ms, 7T with SC72 gradient sets. Full protocol details are available here: https://github.com/layerfMRI/Sequence_Github/blob/master/Terra_protocolls/Fuzzy_ripples/CMRR_ax_1156_slab.pdf

### 3.5) Comparing Fuzzy Ripples across echo spacings and sequences

Six participants were examined to compare Fuzzy Ripples across various echo spacings in popular layer-fMRI sequences: 3D-EPI^12^, CMRR Multiband 2D-EPI^8^, SIEMENS SMS 2D-EPI, and Dual Polarity GRAPPA (DPG) WIP 1105D^16^. These experiments were conducted on SIEMENS 7T Terra scanners. Six sessions were carried out on the 7T Terra scanner at NIMH, with two participants additionally scanned on the NMRF Terra. Parameters such as resolution, TR, TE, acceleration, and FOV were matched across echo spacings, with axial slabs covering the temporal lobes using 36 slices. Echo spacings varied between 1 ms and 1.26 ms. Full protocol details are available here: https://github.com/layerfMRI/Sequence_Github/blob/master/Terra_protocolls/3rd_order_shim/20240528_thirdordershim_siemens.pdf.

### 3.6) Sequence comparison with DPG data

We conducted experiments with two participants to demonstrate the differences between the proposed dual-polarity averaging and the previously proposed dual-polarity GRAPPA (DPG) method^16^. In the DPG approach, differences between read directions across odd and even EPI lines are corrected using a GRAPPA method, where GRAPPA reference data are acquired with two polarities to train separate GRAPPA kernels for odd and even lines. This approach addresses higher-order phase differences more effectively than conventional EPI corrections, particularly mitigating edge ghosts and B0-related artifacts. However, DPG does not target trajectory errors in the readout-direction, which can lead to low spatial frequency artifacts.

Due to the lack of publicly available reconstruction code for DPG, we compared our dual-polarity averaging implementation in 3D-EPI, with the DPG implementation of the SIEMENS WIP 1105 (VE12U) 2D-SMS EPI. We matched echo spacing, resolution, acceleration, TE, and FOV across both sequences. Note that the sequences might use different reconstruction pipelines. Full protocol details are available here: https://github.com/layerfMRI/Sequence_Github/blob/master/Terra_protocolls/Fuzzy_ripples/20230706_segmentationVSGRAPPA.pdf

### 3.7) Imaging the Amygdala and the Cerebellum

Six participants were scanned to evaluate the feasibility of imaging small brain areas at 0.8mm resolution using dual-polarity averaging. Scanning was conducted with 0.8 mm isotropic resolution, partial Fourier 6/8, GRAPPA 3, and 32-channel Rx Nova coils, with a total acquisition time of 14 minutes per functional experiment on a SIEMENS 7T Terra scanner equipped with SC72 gradient sets. Full protocol details are available here: https://github.com/layerfMRI/Sequence_Github/tree/master/low_brain.

Functional tasks included 14 repetitions of 30-second blocks of activation and rest. The amygdala was activated by presenting fearful faces versus objects, and the cerebellum was activated using finger tapping tasks. Results are presented in Fig. 8C and Fig. S5C.

### 3.8) Experiments to exemplify limits of sampling speed at 0.6 mm resolutions

To obtain high-resolution fMRI connectivity datasets with whole-brain coverage, we used the Next Gen 7T scanner^17^ (Feinbergatron, Terra-impulse edition) with its advanced Impulse gradient system (Siemens) featuring a slew rate of 900 T/m/s and a maximum gradient of 200 mT/m. The experiments employed a 64-channel Rx, 8-channel Tx coil (MR CoilTec)^18^. Three participants were scanned at 0.64 mm resolution with 3D-EPI. To cover the whole brain with 180 slices in a reasonable 11-second acquisition time, we used aggressive GRAPPA acceleration (3 x 2). In the low-SNR regime of such small volume voxels, such high acceleration GRAPPA unaliasing is limited by artifacts from EPI phase inconsistencies. We investigated whether dual-polarity averaging can mitigate these artifacts. Additional imaging parameters included: multi-shot 2 segmentation in the in-plane axis, TE = 20 ms, echo spacing = 0.69 ms, BW = 1592 Hz, FOV = 200 x 200 mm, matrix size 314 x 314. Full protocol details are available here: https://github.com/layerfMRI/Sequence_Github/blob/master/dual-polarity/FeinbergatronWholeBrain_invivo20220710.pdf. We utilized 15 minutes of movie clips from the 7T HCP study (MOVIE1) to explore advanced fMRI methodologies.

### 3.9) Functional acquisition contrast

For functional runs, we combined the sequence with VASO imaging. This involved a global inversion pulse that saturates the blood signal every other TR to localize fMRI signal changes without contamination from large draining veins^19^.

### 3.10) Analysis: Image reconstruction, motion correction, dual-polarity averaging, and GLM

MRI data were reconstructed on the scanner^20^ using MOSAIC^21^. Dual-polarity averaging of complex-valued images was performed offline between odd and even TRs following motion correction to allow analysis of both averaged and raw images. For future applications, complex-valued averaging can also be performed in SIEMENS ICE using “short-term” inner-loop averaging between odd and even TRs. This averaging occurs in projection space (post-read FFT but pre-phase FFT) before EPI phase correction on a coil-by-coil basis, prior to GRAPPA reconstruction. Note that, apart from potential noise differences, dual-polarity averaging should be effective either way, as GRAPPA and phase-preserving coil combination performed in this work are linear operations.

Motion correction was performed in AFNI (version AFNI_23.2.04) with 3dAllineate. BOLD correction of VASO data was performed in LayNii^22^ (version v2.7.0).

Auditory fMRI data were denoised with NORDIC^23^ as described by Knusden et al.^24^. Functional activation analyses were performed with GLM implementation of AFNI’s 3dDeconvolve. Layerification was performed in LayNii with the equi-volume principle in LN2_LAYERS.

All analysis scripts are available on Github: https://github.com/layerfMRI/repository. This included a full script of the preprocessing including complex-valued motion correction and subsequent dual-polarity averaging is available on github: https://github.com/layerfMRI/repository/tree/master/Fuzzy_Ripples.

## 4.) Results

### 4.1) Characterization of Fuzzy Ripples Compared to other EPI ghosts

According to the theory of ramp-sampling EPI, as summarized in Fig. 3, Fuzzy Ripples can be described as a result from gradient trajectory imperfections, distinct from off-resonance-induced Nyquist ghosts. Fig. 4 and Fig. S1 highlight the different spatial characteristics of Fuzzy Ripples compared to conventional edge ghosts. It is visible that Fuzzy Ripples are significantly reduced when EPI trajectories do not use ramp sampling (see panels 4A and 4D). In such cases, eddy currents are expected to have largely decayed by the time k_x_-coordinates with high signal power are acquired, leading to edge ghosting being the primary source of artifacts (panel 4D).

**Fig 4:**
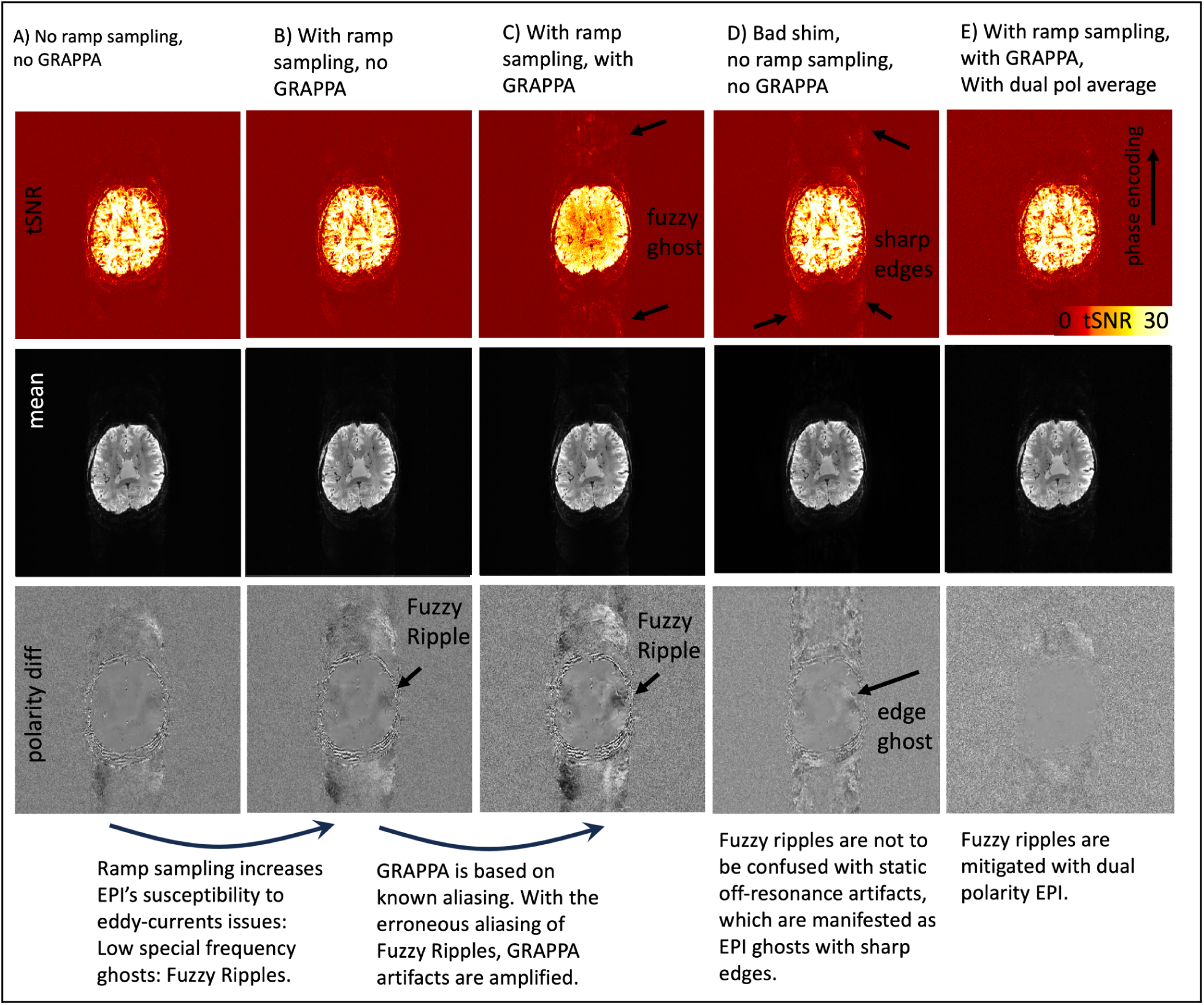
Interaction of Fuzzy Ripples with other common EPI artifacts: GRAPPA ghosts and static off-resonance ghosts. This figure illustrates EPI acquisitions with different combinations of ramp sampling, poor B0 shim, and GRAPPA. The unusually large field of view (FOV) was purposefully chosen to detect peripheral ghost artifacts. Signal differences between reverse EPI polarity images are shown to emphasize spatial ghost patterns that might be too subtle to observe with conventional image intensity windowing. The read direction in left-right, phase encoding direction is anterior-posterior. **A)** Without ramp sampling, imaging data are acquired only during the flat top of the gradient waveform. This minimizes the impact of large gradient errors, resulting in relatively weak Fuzzy Ripples in the EPI images. **B)** With ramp sampling enabled, EPI becomes more sensitive to the largest gradient errors, causing Fuzzy Ripples to intensify. These ripples appear as aliasing of low spatial frequencies, with no sharp edges evident in the phase encoding direction. **C)** GRAPPA, which relies on a known aliasing pattern, is affected by erroneous Fuzzy Ripple ghosts and thus amplifies their impact. **D)** This differs from static off-resonance effects. For instance, with suboptimal shimming (deliberately altered in this case), the off-resonance effects do not amplify the low-spatial frequency Fuzzy Ripples. Instead, they introduce edge ghosts at high spatial frequencies, which differ from Fuzzy Ripples in their appearance. **E)** The dual-polarity averaging approach effectively mitigates both sources of artifacts. The resulting images are nearly artifact-free. Acquisition parameters of data presented here are mentioned in methods section 3.3. See supplementary Figure S1 for the reproducibility of these results in another participant and on another scanner.

Conversely, when ramp sampling is employed, along with the associated trajectory imperfections closer to the center of k-space, Fuzzy Ripples become more pronounced (panel 4C). These aliasing patterns are further exacerbated when GRAPPA is used. However, dual-polarity averaging effectively mitigates Fuzzy Ripples (panel 4E).

The findings depicted in Fig. 4 and Fig. S1 support the idea that Fuzzy Ripples arise from trajectory imperfections linked to eddy currents, rather than from conventional B_0_-related off-resonance effects. However, these results do not provide insights into the specific origins of these eddy currents.

### 4.2) Origin of readout-specific eddy currents

Our investigations, informed by findings from Boulant et al.^14^ on a 11.7T scanner, suggest that inductive coupling between 3rd order shim coils and gradient systems can produce significant eddy currents at certain switching rates. Even when EPI Fuzzy Ripple ghosts are relatively mild, respiration-induced B0 fluctuations during fMRI can cause alternating constructive and destructive interference between the main signal and the ghost, potentially reducing fMRI stability and detection sensitivity.

To explore the impact of 3rd order shim coils on high-resolution fMRI stability, we compared task-based activation maps obtained with and without the 3rd order shims. As shown in Fig. 5 and Fig. S2, Fuzzy Ripples (indicated by white arrows) are present in areas where there is minimal significant fMRI activity. Unplugging the 3rd order shims reduced this artifact, demonstrating their role in the observed distortions.

**Fig 5:**
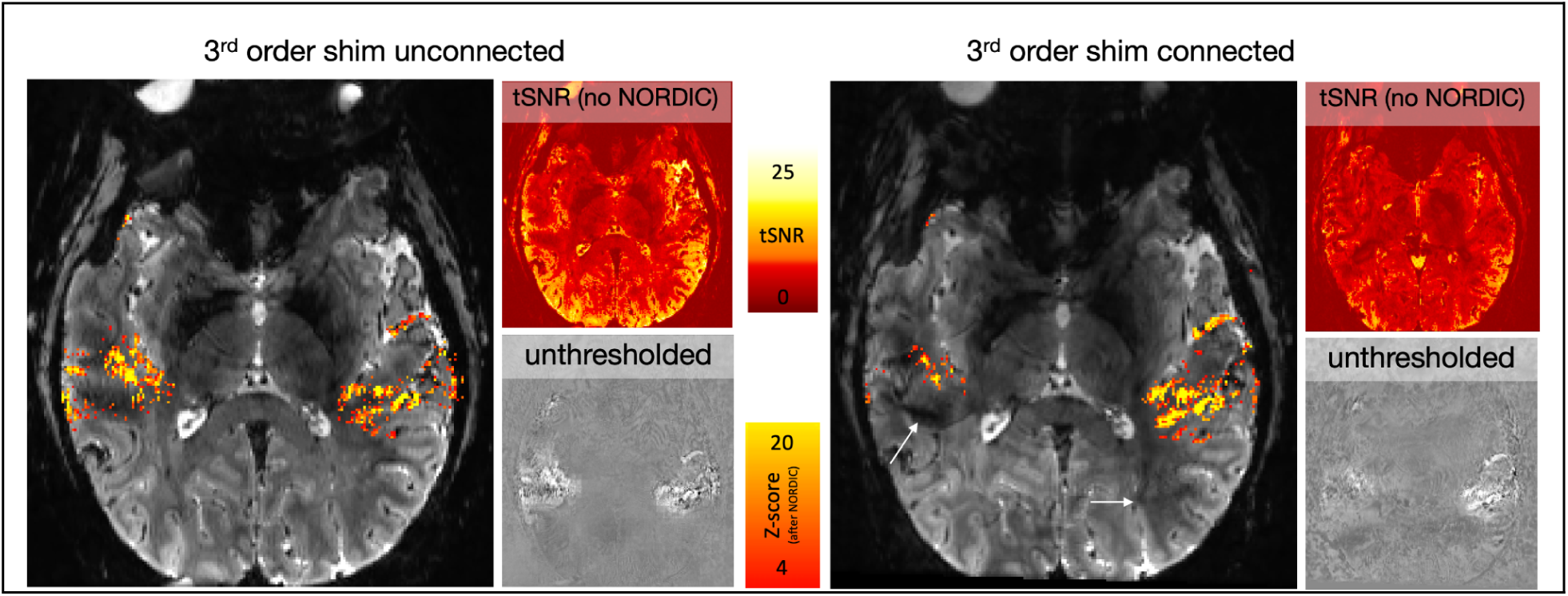
Impact of 3rd order shim-induced Fuzzy Ripples on fMRI activation detectability. This figure demonstrates how Fuzzy Ripples, induced by the 3rd order shim, can affect the detectability of fMRI activation. These data refer to 2D-EPI with block designed auditory activation with NORDIC denoising. When the 3rd order shim is connected, the Fuzzy Ripples can be so pronounced that they mask parts of the auditory activation, preventing it from reaching the detection threshold. White arrows indicate areas where the Fuzzy Ripples are more intense with the 3rd order shim engaged. Although Fuzzy Ripples are still present when the 3rd order shim is disconnected, they are less severe. Acquisition parameters of data presented here are mentioned in methods section 3.4. See Supplementary Fig. S2 for a replication of these findings in a different participant.

### 4.3) Mitigation strategies of Fuzzy Ripples across echo spacings

The results shown in Figures 4 and 5 indicate that Fuzzy ripples can be mitigated through dual-polarity averaging and by disabling the third-order shim, respectively. To evaluate the effectiveness of these mitigation strategies across a broader range of potential fMRI acquisition protocols, we tested them over various EPI echo spacings. The results, presented in Figures 6 and S3, demonstrate the outcomes of these experiments. It is visible that Fuzzy ripples are most pronounced at echo spacings around 1.26 ms and 1 ms. The latter is expected, as it is near the ‘forbidden frequencies’ associated with known mechanical resonances of the SC72 gradient set. The 1.26 ms echo spacing may be related to the fact that the third harmonic of the EPI wave form (1190 Hz) is overlapping with a mechanical resonance of the x/y direction of the SC72 gradient (1100 Hz ± 150 Hz). The results show that these Fuzzy ripples are mitigated when the third-order shims are unplugged (Figure 6B). Additionally, even with the third-order shims enabled, dual-polarity averaging can reduce the Fuzzy ripples at the most problematic echo spacing of 1.26 ms (Figure 6C).

**Fig 6:**
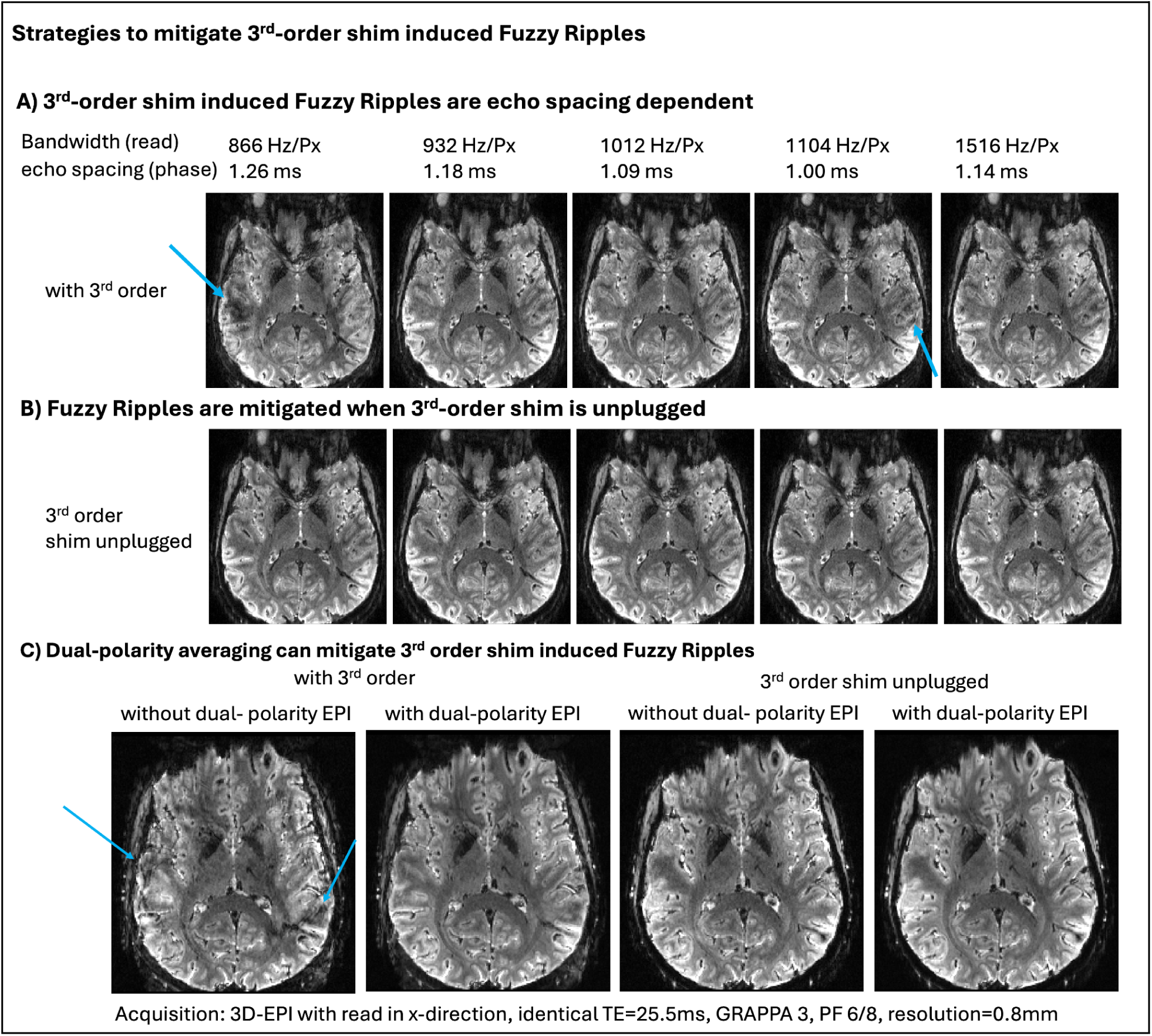
3^rd^-order shim-induced Fuzzy Ripples as a function of echo spacing, dual-polarity averaging, 3rd order shim. **A)** The Fuzzy Ripple artifact varies with the echo spacing of the EPI readout. Consequently, the strength of this artifact can be reduced by adjusting the readout protocol, although such adjustments may compromise TE and readout efficiency. **B)** The Fuzzy Ripples induced by the third-order shim can be mitigated by disconnecting its circuit. Opening this circuit reduces the inductive coupling between the third-order shim and the gradient and reduces Fuzzy Ripple artifacts. **C)** As indicated by Figures 3 and 4, dual polarity averaging can counteract the Fuzzy Ripples induced by the third-order shim. This approach can mitigate fuzzy ripples, even for the most problematic echo spacings with the third-order shim still connected. Though, faint residual fuzzy ripples remain. Acquisition parameters of data presented here are mentioned in methods section 3.5. Supplementary Figure S3 presents a reproduction of the results shown here.

### 4.4) Comparison with other dual-polarity approaches

We evaluated the efficiency of dual polarity averaging in comparison to other methods promoted for mitigating EPI ghosting in high-resolution UHF protocols. These tests were performed with the third-order shim connected and an echo spacing of 1.01 ms. Such protocols are commonly used in layer-fMRI because they enable the fastest acquisition times, shortest TE, and matrix sizes of 250-300.

Panel 7A shows a 2D-EPI image using the CMRR multiband sequence, where Fuzzy Ripples (highlighted by green ellipses) and off-resonance effects (highlighted by yellow ellipses) are visible. Panel 7B illustrates the use of the dual polarity GRAPPA sequence, as conceptualized by Hoge et al^16^ and distributed for SIEMENS VE as part of the WIP package 1105. This approach reduces off-resonance artifacts, but some Fuzzy Ripples remain, appearing as fuzzy dark shading patterns (green ellipses). A k_x_-specific dual-polarity GRAPPA kernel may mitigate these residual Fuzzy Ripples more effectively^24^ (Wang, 2024).

However, the DPG data show a lower temporal signal-to-noise ratio (tSNR) compared to single polarity data. This reduction in tSNR has previously been hypothesized to be related to the ‘Sodickson paradox,’ which occurs when using larger GRAPPA kernel sizes and less GRAPPA fit regularization. With publicly available reconstruction code that allows for the optimization of GRAPPA parameters, this tSNR reduction might be addressable.

For 3D-EPI, which we employed in the dual-polarity averaging approach (panels 7C and 7D), we observed that Fuzzy Ripples can be more effectively mitigated.

### 4.5) Applications of dual-polarity EPI in protocols challenged by Fuzzy Ripples

The results presented in Figs. 5–7 demonstrate that the proposed mitigation strategies, including dual-polarity averaging and disconnecting third-order shims, can effectively reduce Fuzzy Ripple artifacts. This suggests the possibility of utilizing layer-fMRI protocols that were previously hindered by Fuzzy Ripples. To illustrate the utility of these mitigation strategies, we applied them to a series of layer-fMRI protocols that were previously unattainable: 1) higher spatial resolution, 2) faster sampling with aggressive acceleration, and 3) targeting lower brain areas.

**Fig 7:**
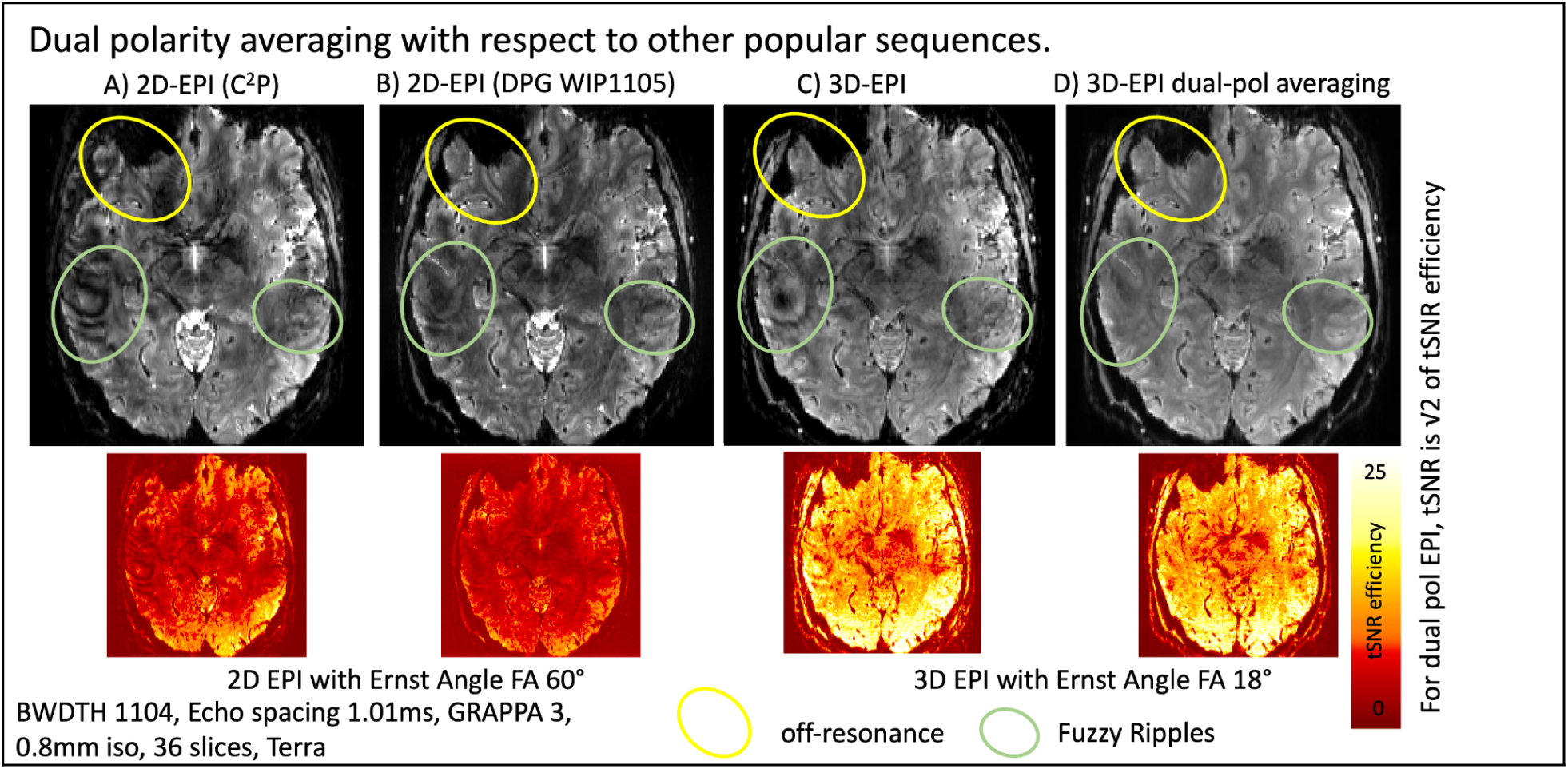
Dual polarity averaging with respect to other popular sequences. All sequences are used with the same resolution, echo spacing, and acceleration parameters. A) This panel shows the CMRR multiband sequence with these protocols, where off-resonance effects and Fuzzy Ripple artifacts are clearly visible. B) This panel displays the MGH simultaneous multi-slice sequence with the option of dual polarity GRAPPA. While off-resonance effects are mitigated, Fuzzy Ripple artifacts, though reduced, remain visible. C) This panel illustrates the same protocols using 3D-EPI. Due to its different Mz steady-state behavior, 3D-EPI inherently has a higher signal-to-noise ratio (SNR). Additionally, off-resonance effects are less noticeable, as they are smeared and partially averaged out. However, 3D-EPI still suffers from Fuzzy Ripples. D) This panel depicts 3D-EPI with dual polarity averaging. It is visible that Fuzzy Ripples are effectively mitigated. Acquisition parameters of data presented here are mentioned in methods section 3.6. Supplementary Figure S4 presents a reproduction of the results shown here.

Figures 8 and S5 present several examples, including 0.65 mm resolution at 3T, whole-brain coverage at 0.64 mm with GRAPPA 2×3, and submillimeter fMRI in the cerebellum, among others.

**Fig. 8:**
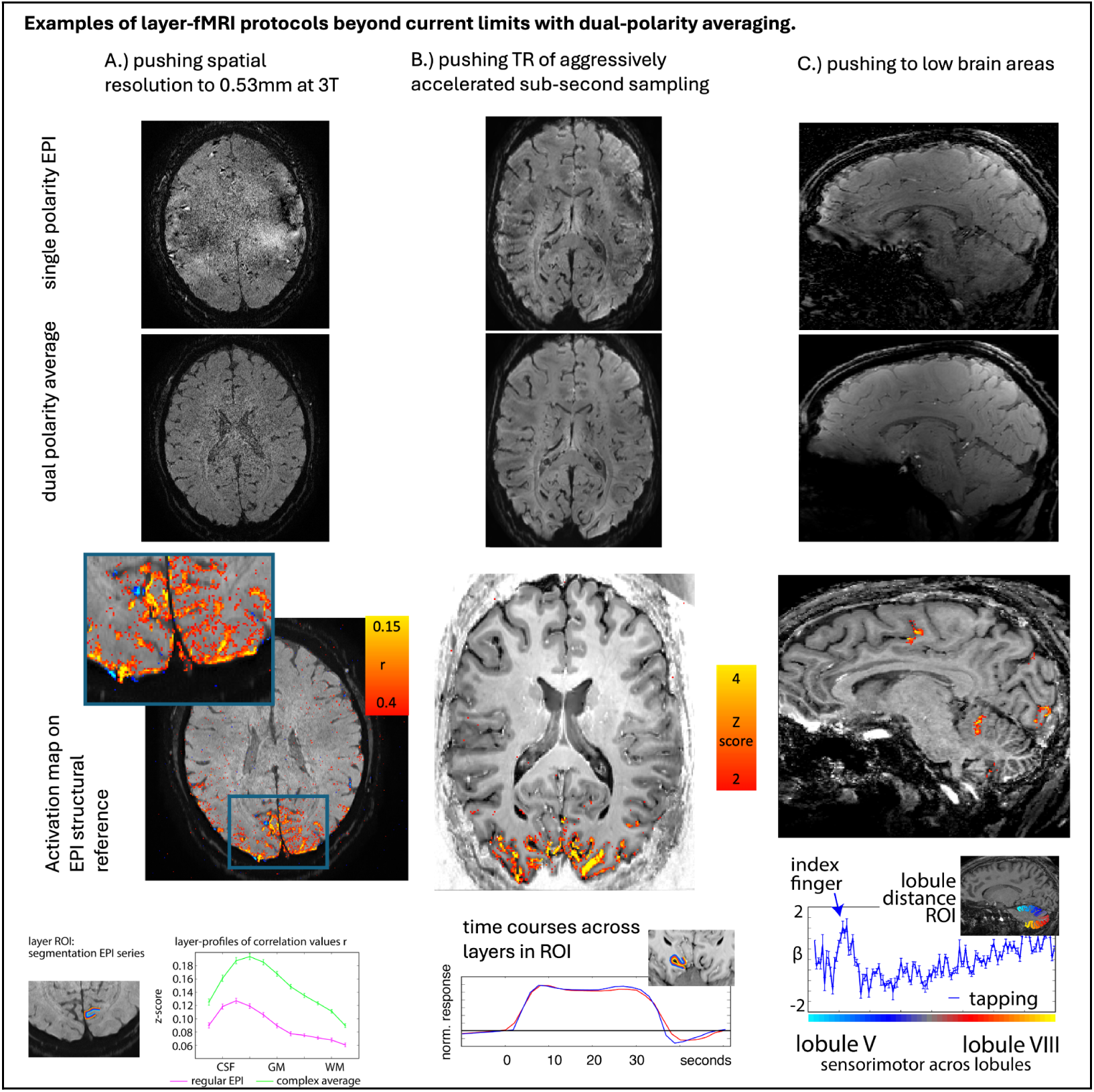
Examples of high resolution protocols pushing the limits of conventional protocols. The individual panels illustrate that achieving high spatial resolution, rapid sampling (through acceleration), and imaging of lower brain areas is challenged by Fuzzy Ripples (as demonstrated in Figure 1). However, dual polarity averaging can mitigate these challenges, enabling the extension beyond the current limitations of conventional layer-fMRI protocols. **A)** 3T Prisma, 3D-EPI with GRAPPA 3, 15-minute movie-watching paradigm, resolution of 0.53 mm. Acquisition parameters of the data shown here are described in methods section 3.2. **B)** 7T Terra, 3D-EPI, GRAPPA 3, three times 14 min checkerboard, resolution 0.82mm, TR_vol_=0.98s for 14 slices. **C)** 7T Terra, 3D-EPI, GRAPPA 3, three averages of a 12-minute finger-tapping experiment, resolution of 0.82 mm. Acquisition parameters of the data shown here are described in methods section 3.7.

## 5) Discussion

In this project, we investigate a prominent artifact of high-resolution Cartesian EPI termed as Fuzzy Ripples - i.e., low spatial frequency signal shadings.

Based on a meta-analysis and experiments involving the gradual increase of spatiotemporal resolution across different brain areas (Figures 1 and 2), we propose that the Fuzzy Ripple artifact is not merely a minor issue in conventional layer-fMRI protocols (0.8 mm, 2-4 s TRs). Instead, Fuzzy Ripples represent a significant source of noise in submillimeter fMRI and pose a greater challenge than thermal noise.

Our empirical studies yielded results that support the hypothesis that Fuzzy Ripples are largely caused by eddy currents in ramp-sampling EPI, which lead to k_x_-specific imperfections in gradient trajectories. In the most advanced SIEMENS whole-body scanners, these eddy currents are largely induced by interactions with the third-order shim. But at a smaller extent, such eddy currents can also be caused by other sources resulting in Fuzzy Ripples in scanners without 3rd order shims. These Fuzzy ripples are exacerbated in the presence of B0 inhomogeneities (as commonly found in lower brain areas), and aggressive GRAPPA accelerations.

We found that the magnitude of Fuzzy Ripples can be reduced using several strategies: 1) disconnecting third-order shims and 2) employing dual-polarity averaging. These approaches enable image acquisition that surpasses the current limitations of resolution, sampling rates, and slice prescriptions.

### 5.1) Significance of exceeding current limits in resolution, sampling rates and flexible slice prescriptions

We have demonstrated that effective strategies can mitigate the Fuzzy Ripple artifact and extend the boundaries of current layer-fMRI protocols. This advancement has significant implications for studying directional neural information flow within and across brain systems in living humans.

#### 5.1.1) Importance of resolution

While conventional layer-fMRI resolutions of 0.8 mm isotropic allow researchers to subsample fMRI activity from different laminar neural populations with varying degrees of partial volume effects, these resolutions represent the bare minimum required. Such resolutions do not permit accurate delineation of structural borders without significant partial voluming, nor do they enable the capture of cytoarchitectonically distinct cortical layers with spatial Nyquist sampling. Achieving a 0.46 mm resolution on conventional 7T scanners (Fig. S5A) would facilitate the direct observation of laminar activation across the cortical ribbon, addressing some of the criticisms faced by the layer-fMRI field^26^. Furthermore, achieving a resolution of 0.53 mm at 3T (Fig. 8A) would help disseminate layer-fMRI beyond the approximately 125 neuroimaging centers worldwide that are equipped with 7T MRI scanners (Huber 2023).

#### 5.1.2) Importance of fast sampling

Recently, 0.8 mm layer-fMRI has been successfully applied with very fast sub-second acquisition windows^27–31^. However, due to the current acquisition constraints related to Fuzzy Ripples, such fast sampling rates have only been possible with small fields of view (FOV). While fast whole-brain imaging is achievable, it is currently limited by Fuzzy Ripple artifacts^26^. The mitigation methods proposed here could make fast whole-brain imaging possible with reduced Fuzzy Ripples. For example, dual-polarity averaging has enabled whole-brain fMRI at 0.6 mm resolutions^17^ and whole-brain quantitative functional T1 mapping^27^.

#### 5.1.3) Importance of lower brain areas

While significant progress has been made in imaging the upper regions of the brain, as evidenced by over 250 published papers, the lower brain areas remain underexplored, hindering the application of whole-brain layer-fMRI. Only 3.4% of these publications (layerfmri.com/papers) have investigated the lower brain areas, preventing layer-fMRI from fulfilling its promise of providing a comprehensive whole-brain functional directional connectome. Numerous important neuroscientific hypotheses involving these lower brain areas remain untested. We are optimistic that the mitigation of Fuzzy Ripple artifacts will pave the way for testing these hypotheses. Some examples include:

- Different layers in the entorhinal cortex, hippocampus, and parahippocampal regions are responsible for memory encoding and retrieval^33–36^.
- Different layers in the fusiform face area (FFA) and parahippocampal place area (PPA) receive feedforward-feedback input for neural representations of faces and places^32, 37–38^.
- Different laminar sub-nuclei of the amygdala are involved in visual perception related to emotional memory versus emotional context^39^.
- Unique lobules of the cerebellum contain sensorimotor digit representations^40^.

### 5.2) Disadvantages of proposed mitigation strategies

Throughout this study, we explored various independent strategies for mitigating Fuzzy Ripple artifacts, with a focus on dual-polarity EPI averaging and disconnecting third-order shims. While these strategies show promise in pushing the boundaries of conventional fMRI protocols, they come with certain compromises.

#### 5.2.1) Unplugging 3rd order shim

Unplugging the 3rd-order shim is a straightforward procedure, supported by vendor-provided workflows (refer to www.layerfmri.com/3rdordershim for workflows applicable to 7T Terra and 7T Plus scanners). However, this procedure requires a system reboot that takes 10-15 minutes, which could result in additional costs due to scanner time budgeting. Moreover, this modification means the scanner is no longer in its FDA-approved configuration. Nevertheless, since layer-fMRI is not an FDA-approved medical procedure to begin with, this is unlikely to present a significant issue. Another drawback of unplugging the 3rd-order shim is that it may leave fine spatial variations in B_0_ inhomogeneities uncorrected.

#### 5.2.2) Complex-valued averaging of dual-polarity EPI

We propose acquiring EPI time series with alternating read polarity, followed by complex-valued averaging of corresponding image pairs. When this averaging is conducted for consecutive pairs of images, the resulting averaged data have half the temporal resolution. When this averaging is rather implemented as a sliding window approach, the number of TRs per run is not reduced. However, this sliding window approach may still lead to temporal blurring of signal fluctuations and decouple the fMRI TR from the effective temporal resolution. However, this limitation can be addressed by refraining from pairwise averaging and instead utilizing alternative reconstruction methods:

1. The dual-polarity approach can be implemented on a calibration-based, run-by-run basis. In this method, phase correction is applied independently to each individual TR without sliding-window averaging, which avoids temporal smoothing but does not account for temporally varying Fuzzy Ripples. The sequence would look slightly different (see Fig. 3C Option 2)). This approach is demonstrated in Steen Moller’s implementation within the CMRR MB sequence^41^.
2. Van der Zwaag et al.^10^ introduced the “CP” approach, where the phase difference of each TR pair is estimated separately and applied as a convolution in the projection space, with alternating signs for odd and even TRs. This method prevents temporal smoothing but requires more substantial changes in image reconstruction.
3. Alternatively, one could forgo pairwise averaging and instead account for Fuzzy Ripples in alternating EPI acquisitions through a regression approach during functional activation analysis (as the artifacts alternate between odd/even TRs). This strategy is only feasible if the stimulation and task design are not locked to odd/even TRs.
4. Alternatively, the measured trajectory imperfections could be used in a one-dimensional non-Cartesian reconstruction model, which could solve the artifact without the need for dual-polarity calibration.

#### 5.2.3) Adjusting echo spacing

During our experiments that were aimed at characterizing the spatial features of Fuzzy Ripples relative to other artifacts, we identified additional scan parameters that may warrant further investigation as potential mitigation strategies for Fuzzy Ripples. These include avoiding ramp sampling (Fig. 4, S1), avoiding GRAPPA (Fig. 4, S1), and adjusting echo spacing (Fig. 6, S3). While ramp sampling and GRAPPA are essential components of modern, efficient fMRI protocols, further exploration of echo spacing adjustments may be justified to quantify its effectiveness as a Fuzzy Ripple mitigation strategy.

Fine-tuning echo spacing is a common optimization step performed during the piloting phase of any layer-fMRI protocol. This process is often undertaken to avoid overlap with mechanical resonances of the main EPI frequency or potential sidebands from non-sinusoidal gradient pulses. This study underscores the importance of such optimizations and suggests that echo spacing should also be optimized specifically for Fuzzy Ripple artifacts. However, this approach is constrained by the desired echo times. For matrix sizes greater than 200, the tradeoff in TE could be as large as 3-12 ms, potentially leading to increased signal decay, blurring, and spatial distortions.

## 6) Conclusion

In this study, we have characterized a significant EPI artifact termed as Fuzzy Ripples, which poses a substantial limitation for laminar imaging.

This low spatial resolution EPI ghosting artifact is caused by trajectory imperfections in ramp sampling EPI, restricting achievable spatial resolution, sampling efficiency, and flexibility of FOV prescriptions. Based on the insights of the origin of this artifact from this study, we proposed several mitigation strategies, including dual-polarity EPI and disconnecting 3rd-order shims. Our findings indicate that these strategies can effectively mitigate Fuzzy Ripple artifacts, thereby extending the capabilities of layer-fMRI acquisition protocols.

Acknowledgements

## 7)

## Ethics

The scanning procedures at 3T have been approved by the Ethics Review Committee for Psychology and Neuroscience (ERCPN) at Maastricht University, following the principles expressed in the Declaration of Helsinki. 7T results were acquired under the NIH-IRB (93-M0170, ClinicalTrials.gov: NCT00001360). We thank Shruti Japee for guidance and support with respect to getting privileges for checking pregnancy tests and IRB.

## Scanning

We thank Kanny Chung and Marcela Montrquin for kind help with respect to participant management. We want to thank the Healthineers Robin Heideman, Reinaldo Gabarron, Sunil Patil, and Bernd Stoeckel for discussions and for sharing their approaches of switching off the 3rd order shim. We thank Samantha Ma for help in acquiring data presented in Fig. S5.

## Advice

We thank Simon Robinson for discussions about most appropriate coil-combination methods in IcePat for appropriate estimation of phase data. We thank SIEMENS Healthineers Bernd Stöckel, Sunil Patil, and Reinaldo Gabarron for advice on the procedure on how to unplug the 3rd order shim. We thank Nadine Gradel for chats about challenges of low brain areas. We thank Federico de Martino for the suggestion to compare the EPI quality of the 3D-EPI sequence with and without dual-polarity averaging with the independent sequence and reconstruction performance of the CMRR multiband sequence.

## 8) Data Availability Statement

- Scanning protocols with complete sets of sequence parameters are available here: https://github.com/layerfMRI/Sequence_Github/tree/master/Terra_protocolls/
- Data on Temporal resolution are available on Openneuro: https://doi.org/10.18112/openneuro.ds003216.v3.0.11
- The LayNii software used here is available on Zenodo: https://zenodo.org/doi/10.5281/zenodo.3514297
- A full script of the preprocessing including complex-valued motion correction and subsequent dual-polarity averaging is available on github: https://github.com/layerfMRI/repository/tree/master/Fuzzy_Ripples
- Raw data of third order shim across echo spacings are here: https://layerfmri.page.link/3rdShim_data
- 3D-EPI sequences (incl. VASO) with optional dual-polarity EPI and on-scanner reconstruction of coil-specific dual-polarity averaging can be accessed via the Siemens C2P exchange platform: For European IP addresses https://webclient.eu.api.teamplay.siemens-healthineers.com/c2p and for US IP addresses https://webclient.us.api.teamplay.siemens-healthineers.com/c2p (v2.0 at time of submission). Search for 3D-EPI (by Stirnberg and Huber) provided by DZNE.
- The auditory data and the data across sequences and across bandwidth are available here: https://zenodo.org/records/13326459

## 9) Diversity statement

Recent work in several fields of science has identified a bias in citation practices such that papers from women and other minorities are under-cited relative to the number of such papers in the field^42^. In the human layer-fMRI community the average of the gender citation bias is 84% male, 16% female (https://layerfmri.com/papers/). We obtained the gender of the first author of each reference. By this measure (and excluding self-citations to all authors of our current paper), our references contain 68% male first and 32% female first. This method is limited in that: (i) names, pronouns, and social media profiles used to construct the databases may not, in every case, be indicative of gender identity, and (ii) it cannot account for intersex, non-binary, or transgender people. We look forward to future work that could help us to better understand how to support equitable practices in science.

## Supplementary information

**Figure S1: reproduced results of Fig. 4.:**
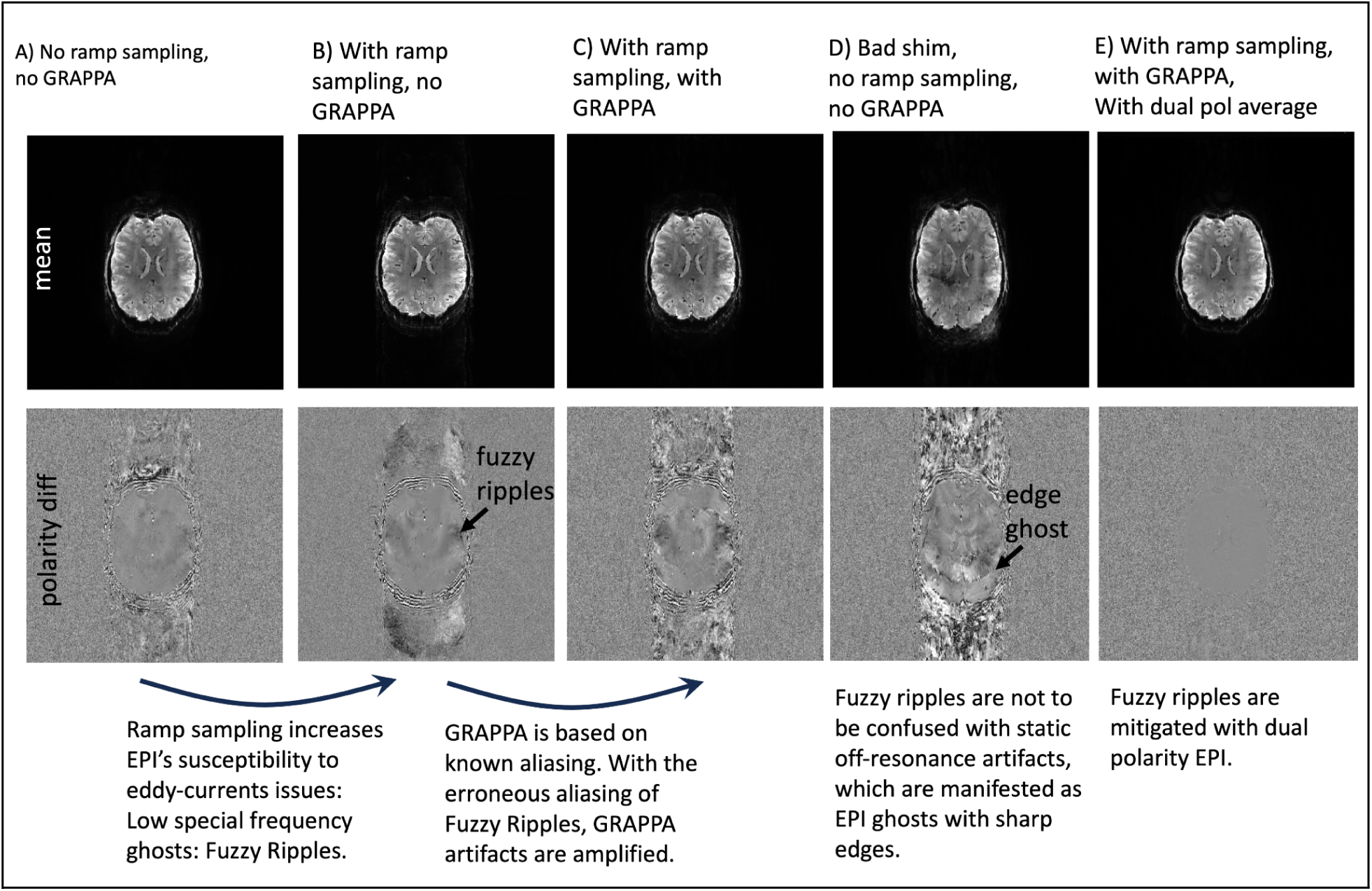
GRAPPA ghosts and static off-resonance ghosts. A series of EPI acquisitions with various combinations of ramp sampling, bad B0 shim and GRAPPA are shown. The FOV purposefully chosen to be unconventionally large, to allow detections of the ghosts in the periphery. The signal difference between reverse EPI polarity images is shown to highlight the spatial ghost pattern that might be too weak to see with conventional image intensity windowing. The read direction in left-right, phase encoding direction is anterior-posterior. **A)** When EPI is done without ramp-sampling, imaging data are solely obtained during the flat top, which eliminates some parts of the largest gradient errors and, thus, the resulting EPI images only show relatively weak Fuzzy Ripples. **B)** When ramp sampling is turned on, EPI becomes additionally sensitive to the largest peaks in gradient errors. Thus the Fuzzy Ripples become stronger. The Fuzzy Ripples manifest as aliasing of low spatial frequencies, as expected. Note that there are no sharp edges in the phase encoding direction. **C)** Since, GRAPPA relies on a known aliasing pattern, which is contaminated with erroneous Fuzzy Ripples, it amplifies the effect. **D)** This is different from static off-resonance effects. For example, in presence of suboptimal shimming (here purposefully altered), it does not amplify the low-spatial frequency fuzzy ripples. Instead, such settings add another source of artifact, namely the edge ghosts at high-spatial frequencies. These sharp borders look different from Fuzzy Ripples. Note that the edge is only sharp along the phase encoding direction. The Fuzzy Ripples that are also amplified with bad shim are smooth in the read directio, **E)** The dual polarity approach can account for both of these sources of artifacts. The resulting images end up almost perfectly flat. Acquisition parameters of data presented here are mentioned in methods section 3.3.

**Figure S2: reproduced results of Fig. 5.**
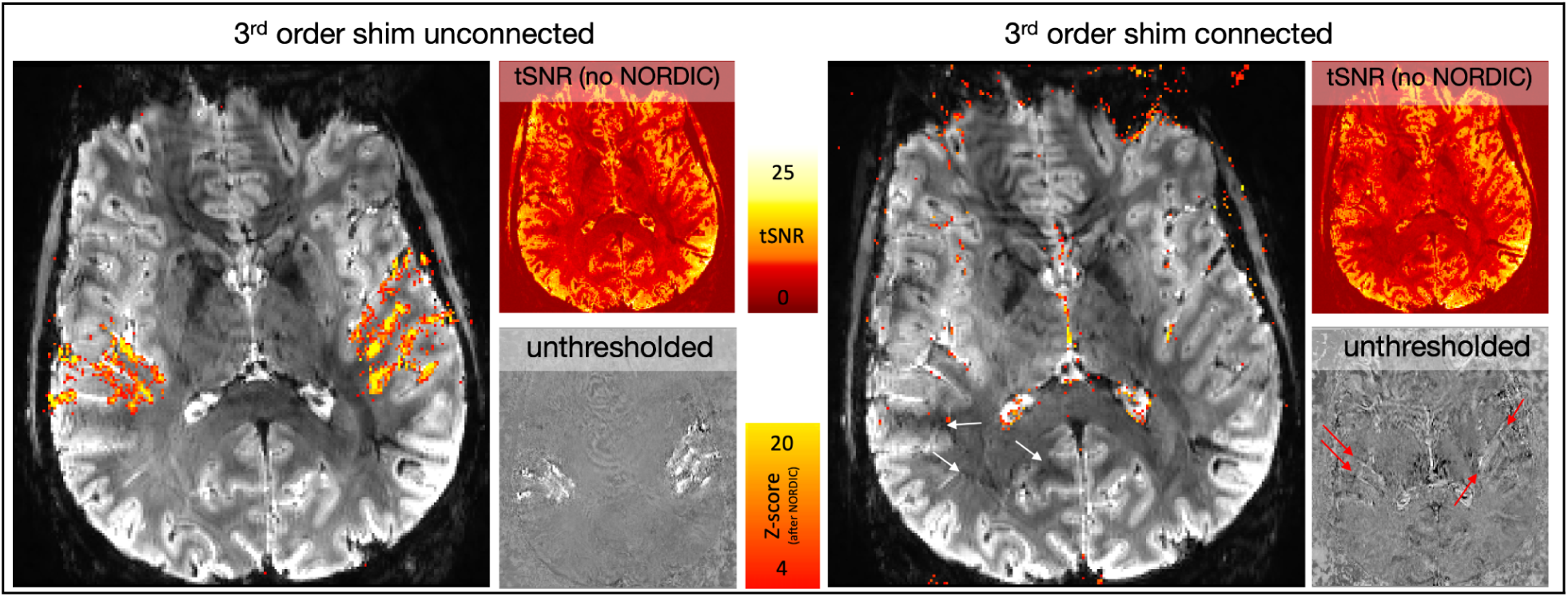
How 3rd order shim induced Fuzzy Ripples affects fMRI activation detectability. When the third order shim is connected, Fuzzy Ripples can be so strong that parts of auditory activation do not exceed the detection threshold. White arrows point to Fuzzy Ripples that are stronger when scanning with the 3rd order shim connected. Fuzzy Ripples are still somewhat present without the 3rd order shim, they are weaker though. Red arrows highlight activated brain areas. They are visible in unthresholded activation maps. However, they are below the detection threshold. Acquisition parameters of data presented here are mentioned in methods section 3.4.

**Fig. S3: reproduced results of Fig. 6.**
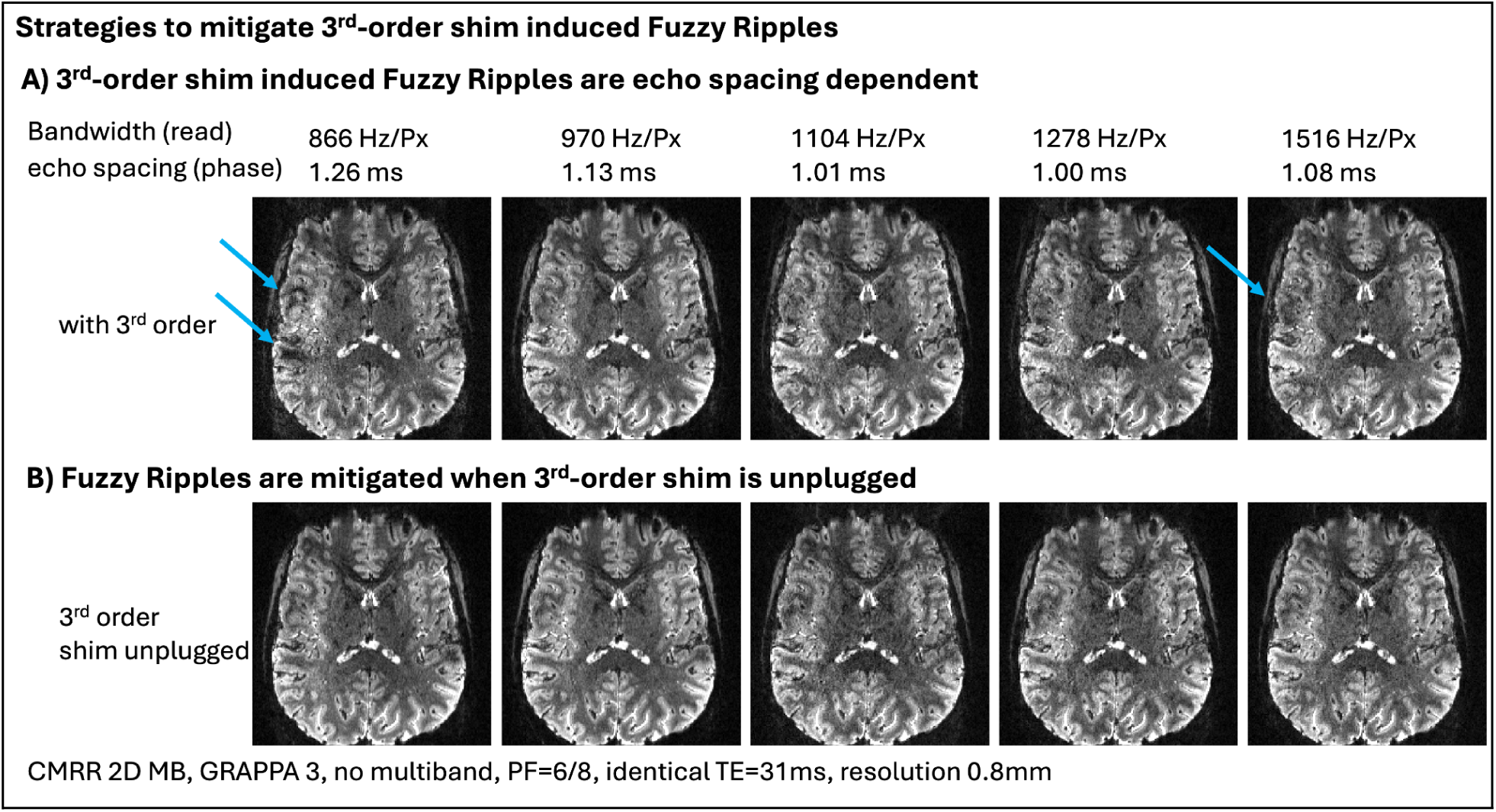
on a different scanner with a different participant using a different EPI sequence: 3^rd^-order shim induced Fuzzy Ripples as a function of echo spacing, cable connected, dual polarity averaging. A) The Fuzzy Ripple artifact is dependent on the echo spacing of the EPI readout. Thus, the artifact strength can be mitigated by protocol adjustments of the readout, which might come along with compromises of TE and readout efficiency. B) 3^rd^-order shim induces Fuzzy Ripples can be mitigated by means of unplugging its circuit. Breaking this circuit reduces the inductive coupling of the 3^rd^-order shim with the gradient. Dual Polarity averaging results shown in Fig. 6C could not be reproduced with the CMRR sequence. Lacking access to the sequence source code, we could not make the required modifications to the sequence. For pilot tests of dual polarity averaging of this sequence in a phantom see (Huber 2023). Acquisition parameters of data presented here are mentioned in methods section 3.5.

**Fig. S4: reproduced results of Fig. 7.**
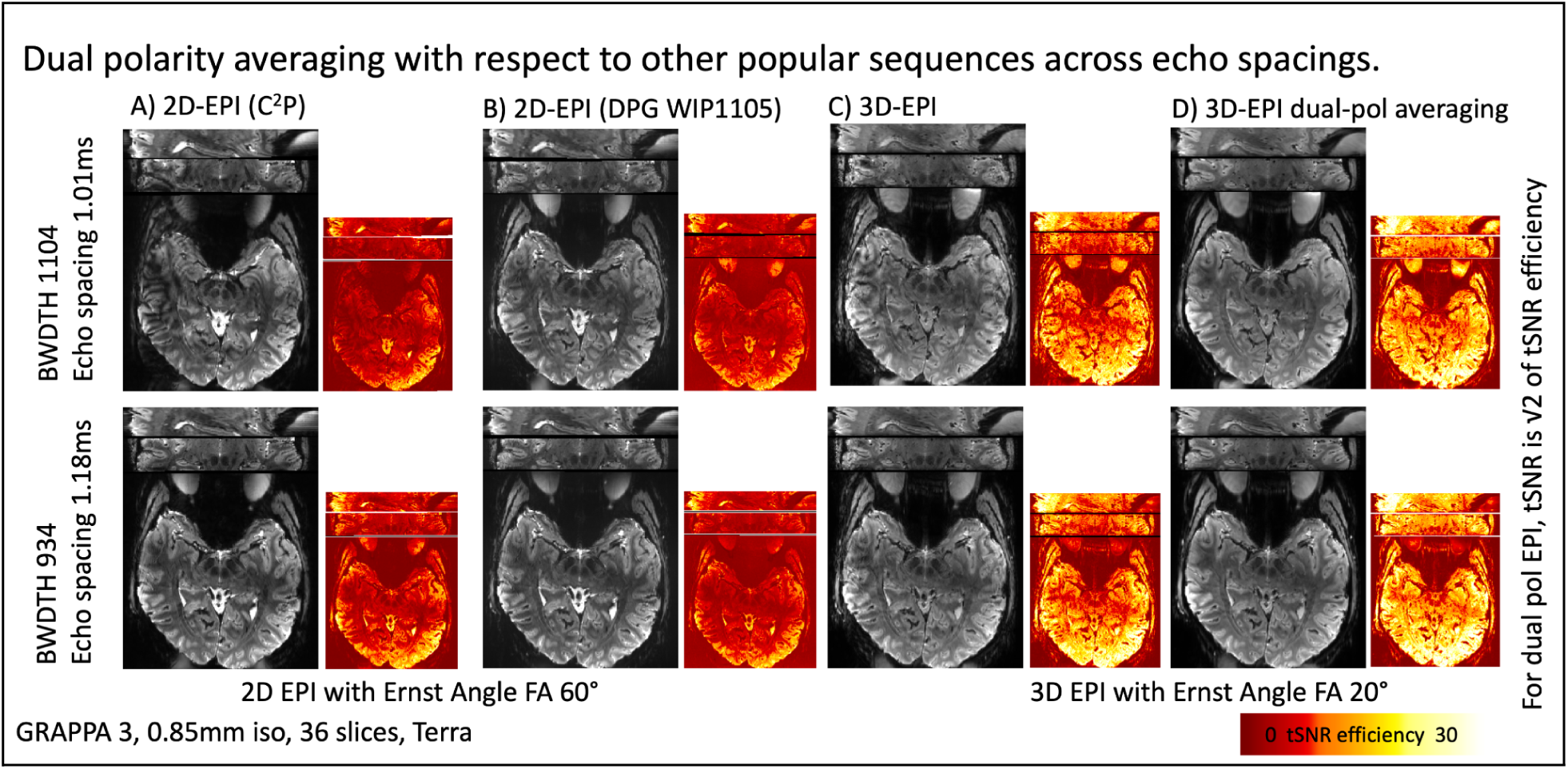
on a different scanner with a different participant with different echo spacings. All sequences are used with the same resolution, and acceleration parameters. Echo spacing is constant for results in each row, respectively. A) depicts the CMRR multiband sequence with these protocols. Fuzzy Ripple artifacts are visible with different strengths across echo spacings. B) depicts the MGH simultaneous multi slice sequence of this protocol with its option of dual polarity GRAPPA. Fuzzy Ripple artifacts in the shorter echos spacing images are mitigated, but still visible. C) depicts the same protocols with 3D-EPI. Due to its different M_z_ steady-state behavior, 3D-EPI has an inherently higher SNR. 3D-EPI suffers from Fuzzy Ripples, especially for shorter echo spacings. D) depicts 3D-EPI with 3D-EPI with dual polarity averaging. It can be seen that Fuzzy Ripples are mitigated across echo spacings. Acquisition parameters of data presented here are mentioned in methods section 3.6.

**Fig. S5: reproduced results of Fig. 8:**
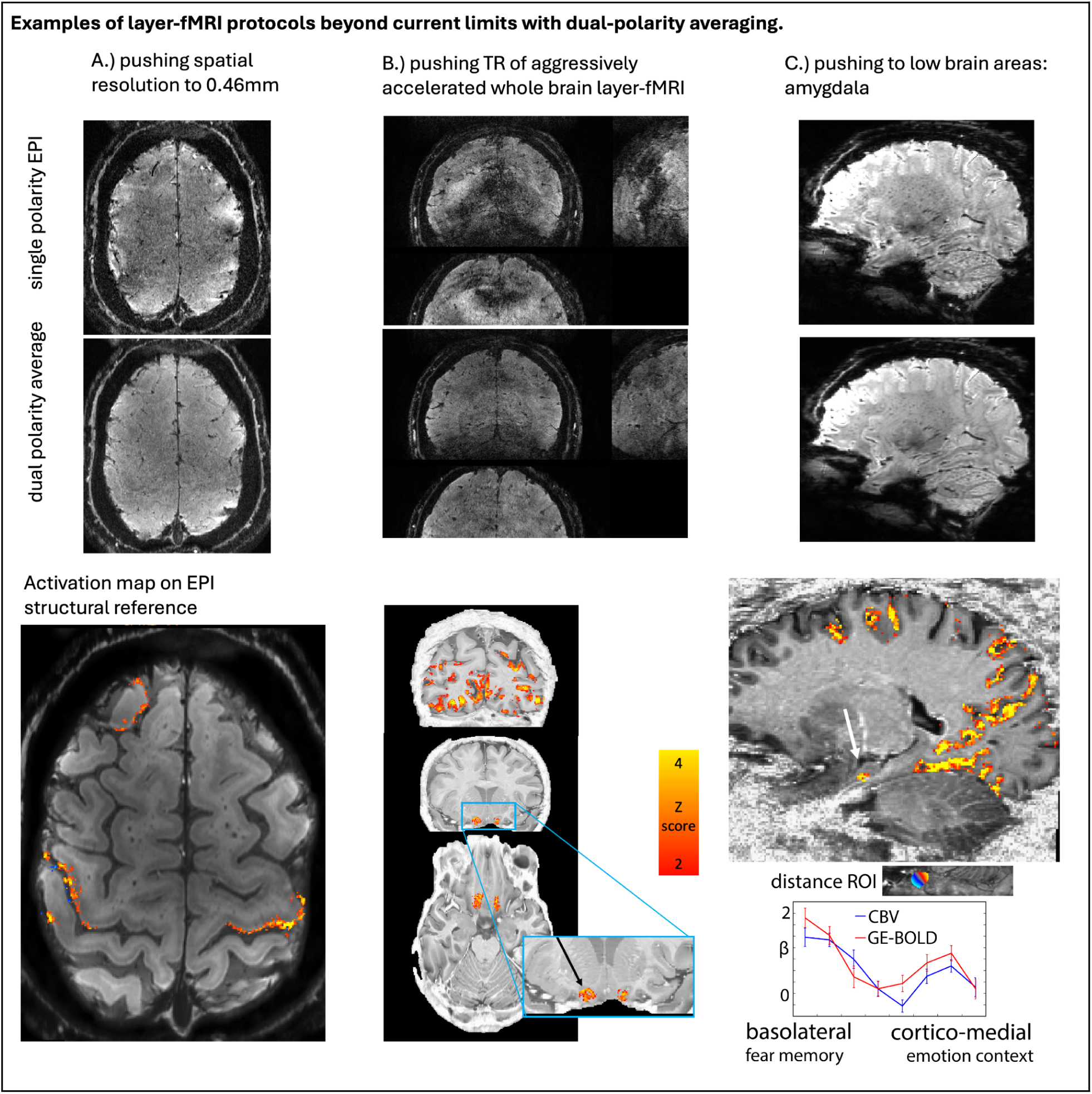
Examples of high resolution protocols pushing the limits of conventional protocols. The individual panels show that pushing to high spatial resolution, to fast sampling (by means of acceleration), and to lower brain areas is challenged by Fuzzy Ripples (as shown in Fig. 1). Dual polarity averaging can mitigate these challenges, thus, allowing to overcome the current limits of conventional layer-fMRI protocols. **A)** 7T Terra, 3D-EPI with GRAPPA 3, 6 fold segmentation, four runs of 12 min finger tapping, resolution of 0.46 mm. **B)** Feinbergatron 7T, 3D-EPI, GRAPPA 8, three 15-minute movie-watching sessions, resolution of 0.64 mm, TR=11.5 s for 314×314×180 voxels. Acquisition parameters of the data shown here are described in methods section 3.8. **C)** 7T Terra, 3D-EPI, GRAPPA 3, three times 15 min emotional faces vs. objects, resolution 0.82mm, Sagittal for deeper brain areas.

